# Predicting Future SARS-CoV-2 Mutations using Deep Learning

**DOI:** 10.1101/2025.07.12.664533

**Authors:** Huzeyfe Ayaz, Ali Reza Ibrahimzada, Arta Armani, Mehmet Baysan, Mehmet Erdem Buyukbingol, Guleser Demir, Elif Karlik, Bulent Ozpolat, Ali Cakmak

## Abstract

SARS-CoV-2 continues to spread over the world steadily as opposed to many earlier estimations that it would disappear in less than two years. Even though SARS-CoV-2 vaccines have reduced the speed of the infection significantly, they could not fully stop it. On the contrary, the World Health Organization has recently published cautionary statements that infection counts are on the rise, and a huge wave is expected in winter. Vaccines mostly target specific regions of the virus. The high mutation rate of SARS-CoV-2 is one essential tool that the virus exploits to escape from the available vaccines. Therefore, researchers have been working on designing next-generation vaccines against the new variants of the virus. Nevertheless, SARS-CoV-2 acquires new mutations faster than we can adapt our vaccines due to long clinical trial periods. Hence, there is a need for computational tools that can predict future SARS-CoV-2 mutations before they even emerge. In this paper, we propose several deep-learning-based methods to estimate the possible future mutations in SARS-CoV-2 genome. We design and evaluate various ensemble and bagging architectures enriched with a large set of genomic, biochemical, and phylogenetic features. We evaluate our models on the GISAID data and demonstrate that the best-performing methods achieve an F1-Macro score of 0.78.

## 1 Introduction

**S**ince the first case is reported in December 2019, 6+ million people have died due to SARS-CoV-2. Following the emergency approval of the first vaccine in December 2020 (by United Kingdom’s Medicines and Healthcare products Regulatory Agency), the death rate of the virus has dramatically decreased. Nevertheless, new life-threatening variants of the virus continue to emerge [1] owing to the high mutation rate of the virus. GISAID [2] reports millions of SARS-CoV-2 variants. Such a high mutation rate quickly deteriorates the effectiveness of the existing vaccines. Therefore, vaccines are now being re-designed to fight against the new variants of the disease. In this race against the virus, scientists have always been many steps behind the virus. This is because the identification of new variants and the adaptation of the existing vaccines to the new variants take a significantly long time. In order to get ahead of the virus, there is a great need for computational tools that can accurately predict the future variants of the virus even before they emerge. This way, scientists may design new vaccines that would work on future variants of the virus as well. In this paper, we propose a computational method to predict the future mutations of SARS-CoV-2. Our method employs k-merized pieces of the SARS-CoV-2 genome along with biochemical and genomic properties of the known mutations as features to build deep learning-based mutation prediction models. Our models also exploit the phylogenetic distances between different strains of SARS-CoV-2. More specifically, we study the prediction performance of three architectures, namely, (i) multi- and single-input ensemble models, (ii) bagging classifiers, and (iii) NLP models for sequence-to-sequence generation. Furthermore, we employ UShER [3] to build a phylogenetic tree of the reported Cov-19 sequences, which contribute features to the constructed prediction models. We evaluate our models on Cov-19 variants compiled by GISAID which contains 2+ million SARS-CoV-2 sequences from all over the world. Our results show that the single-input ensemble model perform the best among the evaluated models yielding an F1-Macro score of 0.78.

Previous works in the literature initially focused on influenza viruses. In particular, predicting the evolutionary behavior of the influenza viruses by estimating which existing strains will dominate the next year’s epidemic has been a popular research focus [4], [5]. Another explored direction was to estimate the influenza strains that could be prevented by vaccines targeting a specific gene of the virus [6], [7]. Besides, predicting dangerous strains [8] and forecasting mutations in influenza [9], [10], [11] were also explored. On the other hand, works on SARS-CoV-2 have a relatively narrower scope and coverage owing to the short history of the virus. Earlier studies focus on the diagnosis of SARS-CoV-2 with AI-based methods [12], [13], [14], estimating how prevalent a strain would be [15], [16], predicting particular mutation rates of nucleotides and aminoacids [17], and genome pattern analysis [18]. Mutation prediction has not been studied in the context of the full SARS-CoV-2 genome. Even though there exist a few studies on influenza mutation prediction, they are not directly applicable to SARS-CoV-2 since (i) influenza genome consists of 8 pieces of strands and even the largest one is much shorter than SARS-CoV-2 genome (i.e., 2.3K vs. 30K), (ii) the mutation rate of SARS-CoV-2 is considerably higher than that of influenza, (iii) the datasets on influenza have been collected for decades, while SARS-CoV-2 datasets are available only for the last 2 years or so, (iv) mutation prediction studies on influenza usually targets only a minor portion of the genome such as a surface binding protein or epitope sites on antigens. Hence, mutation prediction on the full SARS-CoV-2 genome is more complex and challenging. In fact, as part of this study, we compare one of the recent techniques [11] employed for influenza mutation prediction to our methods, and empirically show that it underperforms on the SARS-CoV-2 genome. With this study, we envision that future SARS-CoV-2 vaccines will be designed in a more data-guided way with a longer time frame of efficacy. In particular, our contributions in this paper are as follows:

- To the best of our knowledge, this study is the first attempt in the literature to estimate the future mutations in the full SARS-CoV-2 genome (Haimed et al. [19] focused on mutation prediction in Orf7a gene which constitutes only 1% of the SARS-CoV-2 genome).
- We propose a k-mer-based data point generation scheme where we consider both mutations as well as conservation of the same nucleotide at a particular position as unique “interesting” events to train our models on.
- Our feature set combines multi-dimensional properties about each “interesting” event that cover genomic, biochemical, and phylogenetic aspects in a novel way.
- The mutation dataset is greatly imbalanced rendering most of the standard predictive models ineffective out of the box. We propose several strategies to deal with the imbalanced nature of the dataset.
- In addition to well-known metrics like F1 score and Area-Under-the-Curve (AUC), we propose a Point-Accepted-Mutation (PAM) Matrix-based [20] metric to evaluate the performance of the prediction models.
- We build our models into a web-based tool to make the mutation prediction capability available to researchers worldwide.

This paper is organized as follows. Section 2 reviews the related work in more detail. In Section 3, we present the architectures of our models and the feature extraction process. Section 4 presents the experimental results on the GISAID dataset. Finally, Section 5 concludes with pointers to future work.

## 2 Related Work

Earlier studies that aim to predict the genomic evolution of viruses date back to the pre-SARS-CoV-2 period [8]. In particular, works on different strains of influenza have pioneered the field. In this section, we first review the related studies on influenza and then discuss the works on SARS-CoV-2.

### 2.1 Works on Influenza

Influenza vaccines are renewed every year, and predicting which strains will dominate next year’s flu season is a significant problem to design the most effective vaccine content [5]. In this area, as a vaccine target, some studies focus on the hemagglutinin (HA) protein which is responsible for binding the virus to the infected cell as a vaccine target [6]. The goal is to predict whether an Influenza type A virus can infect human hosts using the HA gene, thus being vulnerable to a vaccine. Moreover, another significant aspect in the development of vaccines and drugs is antigenic variation. Although there have been promising studies on modeling antigenic variation using amino acid sequences, studies focusing on deep learning are demonstrated to be more appropriate for the task. As an example, Xia et al. [7] aim to predict the antigenic variation of influenza A H3N2 by incorporating amino acid changes and other features that are affiliated with antigenicity into a convolutional neural network (CNN) and bidirectional long-short-term memory (BLSTM). Another study [4] uses sampled phylogenetic trees based on HA sequences from human influenza viruses to predict which influenza virus strains are likely to be successful. Iwasaki et al. [8] take a different direction on predicting hazardous strains. The authors attempt to predict the directionality changes of virus sequences with the assumption that such re-arrangements may increase the harm potential of a virus strain.

Finally, mutation prediction on influenza viruses is another concentration area in the field. LSTM networks are widely used in sequence prediction because of their ability to learn long-term dependencies. For example, Mohamed et al. [9] conducted one of the studies using the LSTM networks. The authors predict the next generation sequences of influenza virus using the popular seq2seq LSTM neural network. In another study, Yan and Wu [10] more traditional methods, i.e., shallow neural networks which demonstrate limited predictive performance. Besides, by using HA sequence data, it is possible to predict whether mutations will likely occur in the next flu season. To this end, Yin et al. [11] propose a recurrent neural network model for mutation prediction in influenza A viruses. However, they predict mutations only in epitome sites which cover around 100+ bases. None of the above listed mutation prediction works on influenza targets the entire virus genome. Instead, they either focus on a particular gene (e.g., HA) or small number of selected sites (e.g., epitomes).

### 2.2 Works on SARS-CoV-2

In this section, we first discuss in general the AI-based prediction and detection regarding different aspects of SARS-CoV-2 and then move to more relevant works.

With the surge of overwhelming numbers of fatal SARS-CoV-2 cases in the early period of the pandemic, the diagnosis of the infection with AI models was one of the hot research areas in the field. By using deep learning models trained on CT scans and X-ray images, SARS-CoV-2 could be diagnosed with high accuracy [12], [13], [14], [21], [22], [23].

Later, the focus shifted more to the analysis of the evolution in the SARS-CoV-2 genome. For instance, several studies track genomic variations in the virus genome [24], [25], [26]. Another relevant problem is to predict the mutation rate of the virus to reveal the frequency of nucleotide and aminoacid replacements in a pairwise manner [17]. To this end, the authors employ a Recurrent Neural Network-based (RNN) Long Short Term Memory (LSTM) model to estimate the evolution of particular mutation rates in future generations of the virus. One limitation of this work is that it considers only single base substitutions. In addition to mutation rates, frequent pattern analysis within the SARS-CoV-2 genome was also studied [18]. The authors employ the existing frequent sequence pattern mining algorithms to first extract sequence patterns and then mine for interesting association rules between frequently occurring nucleotide sequences. Moreover, they also train probabilistic sequential models, such as Hidden Markov Models, to predict a nucleotide at a particular position based on the nucleotide sequence located prior to the position of interest in the SARS-CoV-2 genome.

Not all SARS-CoV-2 mutations are interesting to researchers. That is, a mutation would be of interest if it would lead to a clinically threatening strain of the virus. As an example, Nagy et al. [16] aim to predict whether a mutation is mild or severe using available clinical follow-up data and nucleic acid sequences downloaded from the GISAID virus repository. They combine a classification algorithm with a feature selection algorithm to link mutation signatures and results. They also developed an online analysis platform where they can predict the clinical outcome, expressed as the probability of severity. Then, Huang et al. [27] extended Nagy et al’s study by considering a larger set of features and classification algorithms. Besides, they also suggested effective classifiers for predicting SARS-CoV-2 patients’ clinical status. Both studies predict the effect of given mutations rather than future possible mutations in the SARS-CoV-2 genome. With a somewhat related goal, Al-Maliki et al. [28] employ classifier models to identify which of the existing variants of SARS-CoV-2 are likely to acquire future mutations. Then, such predictions are exploited as a proxy to estimate the variants with higher potential to spread, i.e., candidates to be strains of interest.

Furthermore, phylogenetic analysis of SARS-CoV-2 strains has been another field of interest. For instance, Haimed et al. [19] proposed a viral reverse engineering approach and studied five viral families, namely, Orthomyxoviridae, Retroviridae, Filoviridae, Flaviviridae, and Coronaviridae. The authors aim to find patterns in genomic sequences and understand how SARS-CoV-2 behaves by comparing different viral families. Finally, they also employ phylogenetic trees of SARS-CoV-2 sequences to predict the next evolved strain by using Long Short-term Memory (LSTM). The last part of this study is closely related to our work. Nevertheless, the major drawback was that they predict mutations in a very small portion of the genome, i.e., ORF7a region which contains only 365 base pairs. Given that the SARS-CoV-2 genome contains around 30,000 base pairs, the scope of Haimed et al.’s study is quite limited.

## 3 Methods

Our mutation prediction pipeline consists of three steps: (i) feature extraction, (ii) data pre-processing, and (iii) model building and evaluation. Feature extraction deals with manipulating the data in different ways and generating new features. In the data pre-processing step, generated features are processed and turned into a suitable format that the models expect in the next step. Moreover, the data is partitioned into different splits such as train, validation, and test. Finally, the last phase involves building several predictive models with different architectures, training, parameter tuning, and performance evaluation steps. We next explain these steps in detail.

### 3.1 Dataset

We obtained SARS-CoV-2 variants from GISAID [2]. The dataset covers all observed mutations between the dates of Dec. 30, 2019, and Oct. 01, 2021. However, the dates of some mutated sequences were missing in our dataset. Therefore, we analyzed the NA ratio of each protein region and removed the sequences with unknown (NA - “not available”) dates from the dataset. The total number of mutated nucleotides per ORF is presented in Table 1. In particular, the table shows that ratios of the NA values are consistent through different ORFs. Therefore removing the sequences which have unknown dates does not affect the balance of the dataset. After the removal of sequences with NA dates, we lost 587,208 mutated nucleotides and the final dataset has 548,128 sequences/nodes. We build [3] a phylogenetic tree of the variants to model the evolution of the SARS-CoV-2 genome. Using the phylogenetic tree, we maintain the evolutionary temporal order of the variants.

**TABLE 1:** Total Mutations and NA ratio for each ORF.

| With NA |  | Without NA |  | NA Rate |
| --- | --- | --- | --- | --- |
| ORF | Total Mutations | ORF | Total Mutations |  |
| S | 236245 | S | 146144 | 0.381 |
| ORF1ab | 951130 | ORF1ab | 588674 | 0.381 |
| ORF8 | 40360 | ORF8 | 24504 | 0.392 |
| N | 115928 | N | 72849 | 0.371 |
| ORF7a | 38364 | ORF7a | 23502 | 0.387 |
| E | 10565 | E | 6810 | 0.355 |
| M | 37858 | M | 23794 | 0.371 |
| ORF3a | 79332 | ORF3a | 48089 | 0.393 |
| ORF7b | 8523 | ORF7b | 5186 | 0.391 |
| ORF10 | 9838 | ORF10 | 6019 | 0.388 |
| ORF6 | 12193 | ORF6 | 7557 | 0.380 |
| **Total** | **1540336** |  | **953128** | **0.381** |

**TABLE 2:** Test results of the model trained with the imbalanced dataset: Performance results with different metrics.

|  | Precision | Recall | F1-Score | Support |
| --- | --- | --- | --- | --- |
| 0 | 1.00 | 1.00 | 1.00 | 12173845 |
| 1 | 0.00 | 0.00 | 0.00 | 395 |
| accuracy |  |  | 1.00 | 12174240 |
| macro avg | 0.50 | 0.50 | 0.50 | 12174240 |
| weighted avg | 1.00 | 1.00 | 1.00 | 12174240 |

PPR: 0.00 NPR: 1.00

**TABLE 3:** Test results of the Model: Performance results with different metrics.

|  | Precision | Recall | F1-Score | Support |
| --- | --- | --- | --- | --- |
| 0 | 0.71 | 0.74 | 0.72 | 39500 |
| 1 | 0.73 | 0.70 | 0.71 | 39500 |
| accuracy |  |  | 0.72 | 79000 |
| macro avg | 0.72 | 0.72 | 0.72 | 79000 |
| weighted avg | 0.72 | 0.72 | 0.72 | 79000 |

PPR: 2.50 NPR: 0.38

**TABLE 4:** Test results of the Multi-Input Ensemble Model: Performance results with different metrics.

|  | Precision | Recall | F1-Score | Support |
| --- | --- | --- | --- | --- |
| 0 | 0.64 | 0.95 | 0.76 | 39500 |
| 1 | 0.90 | 0.46 | 0.61 | 39500 |
| accuracy |  |  | 0.71 | 79000 |
| macro avg | 0.77 | 0.70 | 0.68 | 79000 |
| weighted avg | 0.77 | 0.70 | 0.68 | 79000 |

PPR: 2.50 NPR: 0.15

**TABLE 5:** Test results of the Single-Input Ensemble Model: Performance results with different metrics.

|  | Precision | Recall | F1-Score | Support |
| --- | --- | --- | --- | --- |
| 0 | 0.85 | 0.67 | 0.75 | 39500 |
| 1 | 0.73 | 0.88 | 0.8 | 39500 |
| accuracy |  |  | 0.77 | 79000 |
| macro avg | 0.79 | 0.78 | 0.78 | 79000 |
| weighted avg | 0.79 | 0.78 | 0.78 | 79000 |

PPR: 4.71 NPR: 0.32

**TABLE 6:** Test results of the Bagging Classifier: Performance results with different metrics.

|  | Precision | Recall | F1-Score | Support |
| --- | --- | --- | --- | --- |
| 0 | 0.70 | 0.70 | 0.70 | 39500 |
| 1 | 0.70 | 0.70 | 0.70 | 39500 |
| accuracy |  |  | 0.70 | 79000 |
| macro avg | 0.70 | 0.70 | 0.70 | 79000 |
| weighted avg | 0.70 | 0.70 | 0.70 | 79000 |

PPR: 2.33 NPR: 0.43

Figure 1 shows the mutation trend over the time for each coding region. Mutation figures are normalized by the lengths of the corresponding protein regions. Since the mutations before Dec. 2020 are not very common, we exclude that part while drawing the figure for visualization purposes.

**Fig. 1:**
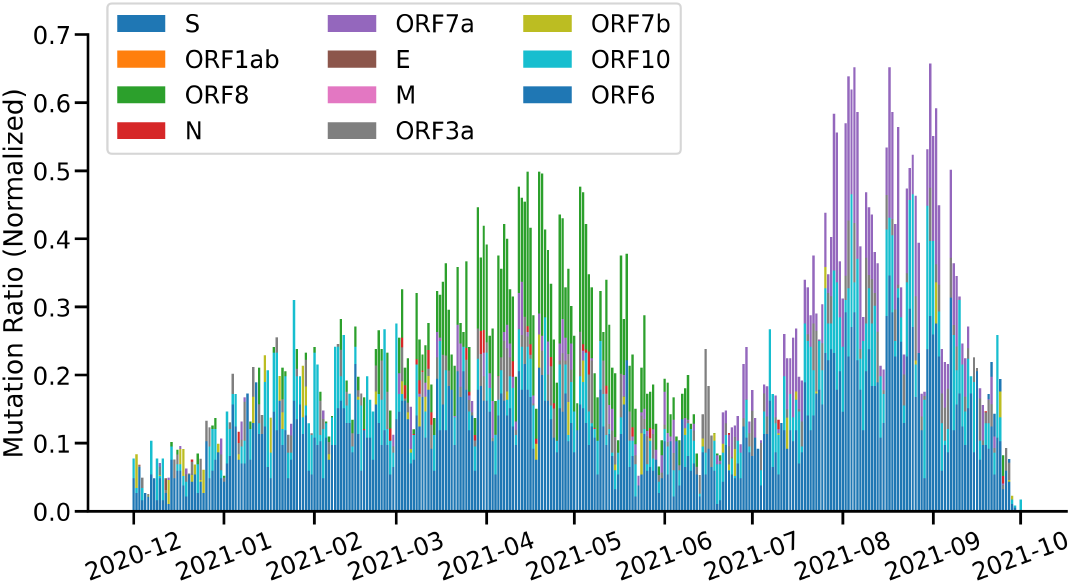
Trend of the mutation ratio for each coding region which is normalized by the corresponding ORF length

The figure shows that mutations are not evenly distributed across different ORFs. As an example, ORF8 and ORF7a regions stand out with higher mutation rates between the dates of Mar. 2021 - Jun. 2021 and Jul. 2021 - Oct. 2021, respectively.

### 3.2 Feature Extraction

Preparing data and extracting features for model training are the key Steps to build effective machine learning models. Using the phylogenetic tree and DNA sequences, we propose a new feature extraction scheme for mutation prediction.

K-mers or k-merization is a well-known method in Bioinformatics to compare sequences and find patterns between them and k denotes a window length as a subsequence [29], [30]. To generate all k-mers for a DNA or RNA sequence, the sliding windows approach is used with one step forward moving, where k is a model parameter to be tuned. However, as different from the standart k-merization process, a residue in position *i* is placed at the center (i.e., midpoint) of *i*th k-mer, i.e., position ⌈*k/*2⌉ in the k-mer. This is to capture the neighborhood of a residue of interest in a k-mer. However, this will lead to blank spaces in the prefix of the first ⌊*k/*2⌋ and the suffix of last ⌊*k/*2⌋ k-mers. Those blank spaces are filled with “-” symbols. As an example, consider the sequence in top left corner of Figure 2 and let k be 5. The first two k-mers would be “- - A T T” and “- A T T C”, respectively, and the last k-mer would be “G G A - -”. Note that for a genome of size *N*, under this scheme, the number of k-mers would be *N*, rather than *N-k+1*.

**Fig. 2:**
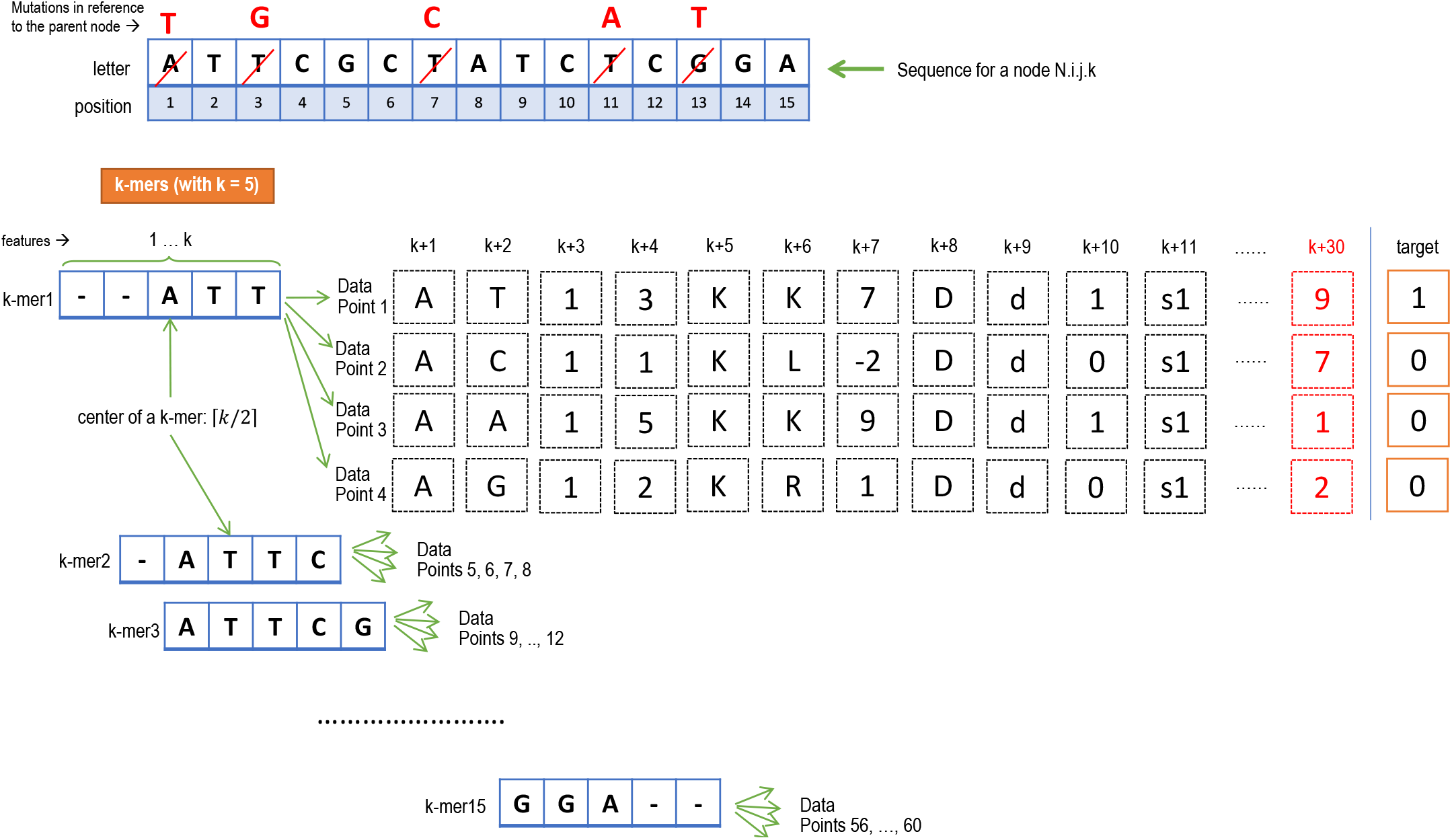
Proposed feature extraction method

For each k-mer, we generate four data points. More specifically, each data point represents a mutation event where the residue at the center of the k-mer is replaced by a different residue (there are 3 possibilities as the DNA alphabet contains 4 letters) or stays the same (i.e., “mutating” to itself or no change). As an example, consider the first k-mer, “- - A T T”, in Figure 2. The position of interest is 1, thus, the center of the k-mer contains the first residue, i.e., “A”. The first data point represents the event that “A” mutates to “T”, the second represents the event that “A” mutates to “C”, the third represents the event that “A” mutates to “A” (i.e., stays the same or no mutation), and finally the fourth represents the event that “A” mutates to “G”.

Each data point is represented as a vector of *k+29* features. We next explain each feature by referring to its position in the feature vector in the following list.

- *1* .. *k*: The first k features contain the nucleotides in the current k-mer. As an example, in Figure 2, the first 5 features of the third k-mer are as follows: [‘A’, ‘T’, ‘T’, ‘C’, ‘G’].
- *k+1*: The original residue at the position of interest in the genome, i.e., the center of the k-mer. As an example, in Figure 2, for the first four data points generated from the first k-mer, *k+1*th feature contains “A”.
- *k+2*: The residue after mutation at the position of interest in the genome. As an example, in Figure 2, for the first data point which represents the mutation event “A” → “T”, *k+2*th feature contains “T”.
- *k+3*: The position of interest in the genome. As an example, in Figure 2, for each of four data points derived from the first k-mer, *k+3*th feature contains value 1.
- *k+4*: Point-Accepted-Mutation (PAM250) [20] score of the mutation at the nucleotide level.
- *k+5*: Amino acid that the nucleotide at position of interest is associated with in the SARS-CoV-2 codon mapping before the mutation.
- *k+6*: Amino acid that the nucleotide at position of interest is associated with in the SARS-CoV-2 codon mapping after the mutation.
- *k+7*: PAM250 [20] score of the mutation at the aminoacid level.
- *k+8*: Number of days that has passed between the emergence of this mutation and the sequencing of the SARS-CoV-2 reference genome.
- *k+9*: Depth of this strain in the SARS-CoV-2 pylogenetic tree that is constructed with the help of UShER [3].
- *k+10*: Indicates whether the mutation is synonymous (1) or non-synoymous (0).
- *k+11*: The name of the ORF that includes the position of interest (if any).
- *k+12* .. *k+29*: Hydrophobicity (k+12 & k+13), polarity (k+14 & k+15), iso-electricity (k+16 & k+17), volume (k+18 & k+19), weight (k+20 & k+21), pKa (k+22 & k+23), pKb (k+24 & k+25), pKx (k+26 & k+27), and pl (k+28 & k+29) of the original AA and the replacement AA, respectively.

Note that each data point represents a mutation event. Hence, if the described mutation in an event is actually observed in the data, it is labeled with value 1 (i.e., target in Figure 2). Otherwise, it is labeled with a target value of 0. As an example, in Figure 2, in position 1, since A has mutated to T, we set the target of the first data point of k-mer1 as 1 (observed mutation in the data) and others as 0.

The features k+8 keeps the value of the estimated elapsed time since the emergence of SARS-CoV-2 till the observation of the current mutation sequence, and k+9 holds the depth of sequence in the phylogenetic tree. In order to estimate the date for an internal node, we employ the following approach.

- Let *θ* be the minimum (*ref_genome_date* − *variant_date*)*/root_to_variant_path_distance* in the phylogenetic tree where reference genome date is 2019-12-01.
- For each internal node X, estimate the date as *minimum*(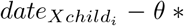 *edge length of child*_*i*_ where *XChild*_*i*_ refers to a child node of X.

*Example:* Consider the phylogenetic tree in Fig. 3 with marked hypothetical edge lengths.

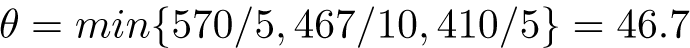

**Fig. 3:**
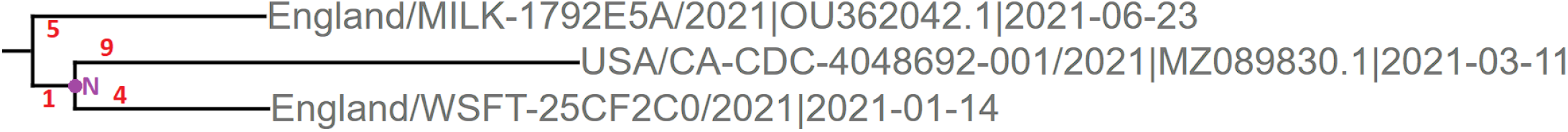
Phylogenetic Tree Example

Estimated date for internal node N:

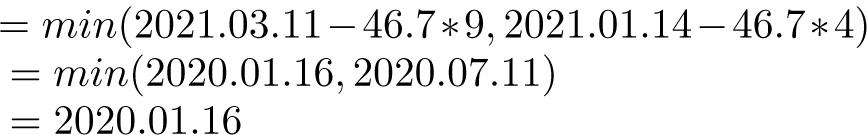

### 3.3 Data Pre-Processing

Applying k-merization to millions of sequences leads to enormous amounts of data. Furthermore, for each k-mer, four data points are generated to cover all possible cases in training. Each data point consists of the combination of categorical (base-pairs, amino acids, etc.) and numerical (position of interest, number of days elapsed, etc.) features. To convert categorical values to machine understandable data, one-hot-encoding is performed. After the initial data generation step, we have roughly 64.5 billion data points each with 205 features which requires roughly 12.6 TB of space on the disk. Training models on such large datasets requires a significant amount of time and computational resources. In order to turn the data into a more manageable form in terms of computational requirements, we sampled the data and randomly selected 117 million data points (around 0.2% of the dataset). During sampling, we preserved the parent-child relationship for the features that need values before and after the mutation.

Since our primary goal is estimating possible future mutations, we choose 2021-08-15 and 2021-09-01 as cut-off dates to split the data into train, validation and test portions. All samples before the 2021-08-15 cutoff date (not-inclusive) are included in the training. All samples remains between the 2021-08-15 (inclusive) and 2021-09-01 (not-inclusive) cutoff dates are included in validation and those that are after the 2021-09-01 (inclusive) cutoff date included in the test split. In particular, train, test, and validation datasets constitute 79%, 9%, and 12% of our sampled dataset, respectively. To preserve the class balance, the splitting into training, validation, and test datasets is done in a stratified manner. Furthermore, within 117 million data points, only 0.0025% of the samples have mutations. Machine learning models do not perform well under such imbalanced class label distributions, and tend to predict all the test samples under the majority label. In order to tackle with this problem, we randomly sampled data points from non-mutated samples of the same size as the mutated samples. The same process is repeated 100 times to gather different samples from non-mutated data points. The main reason for randomly sampling the non-mutated data multiple times is training and evaluating models with non-mutated data as much as possible. Next, we trained separate models using each sampled data separately. In addition, as another direction, we combined all sampled datasets to create a relatively large dataset to train a single model as explained in section 3.4.3.

### 3.4 Model Training & Evaluation

In comparison to the well-known machine learning algorithms like Support Vector Machine (SVM) [31], Logistic Regression [32], Random Forest [33], etc., deep neural networks (a.k.a. deep learning) [34] allow us to create task-specific models with various hyper-parameter options. Moreover, since they consist of many layers and units, they are more capable of extracting patterns from data. Therefore, we study various ways of constructing deep learning models with different architectures. Moreover, a sequence-to-sequence (seq2seq) model called “t5-base”, which is publicly available on the HuggingFace [35] platform is also employed to generate a child mutated genome from a given parent genome.

For each neural network model that we have created from scratch, Tensorflow/Keras [36] library is used. Additionally, Relu [37] activation function is used for each intermediate layers and the Sigmoid [38] activation function is used for the output layer. We employ Adam [39] as an optimizer and binary cross-entropy as a loss function. As for the NLP model, we employ the PyTorch [40] implementation of the models on the HuggingFace platform. When training a model in TensorFlow, the layer naming convention is designed to ensure uniqueness across all layers in the entire model. The “ensemble 5” prefix indicates the 5th member of the ensemble model and each member is treated as a separate submodel. In addition to that, layer name with the occurrence number of this layer is added as a suffix to ensure uniqueness. For example, “batch normalization” is the base name of the BatchNormalization layer, as the name implies. The number placed after the base layer name indicates the n’th occurrence of the layer within the entire model. However, since we trained multiple models in one training session, the numbers may not indicate the exact submodel or layer numbers. We next explain model architectures in detail.

#### 3.4.1 Building a Model with Imbalanced Data

As a baseline approach, we built a model on the original imbalanced dataset as a baseline approach. In particular, we created a simple model that has two fully connected layers (Figure 4). The intermediate “dense” layer downsizes 205 features to 128. Then, the new 128 units are connected to a single unit layer to predict the final label as either 1 (mutated) or 0 (non-mutated). As discussed above, Relu is used as an activation function for the intermediate layers and Sigmoid is used for the output layer. Additionally, Adam is used as an optimizer.

**Fig. 4:**
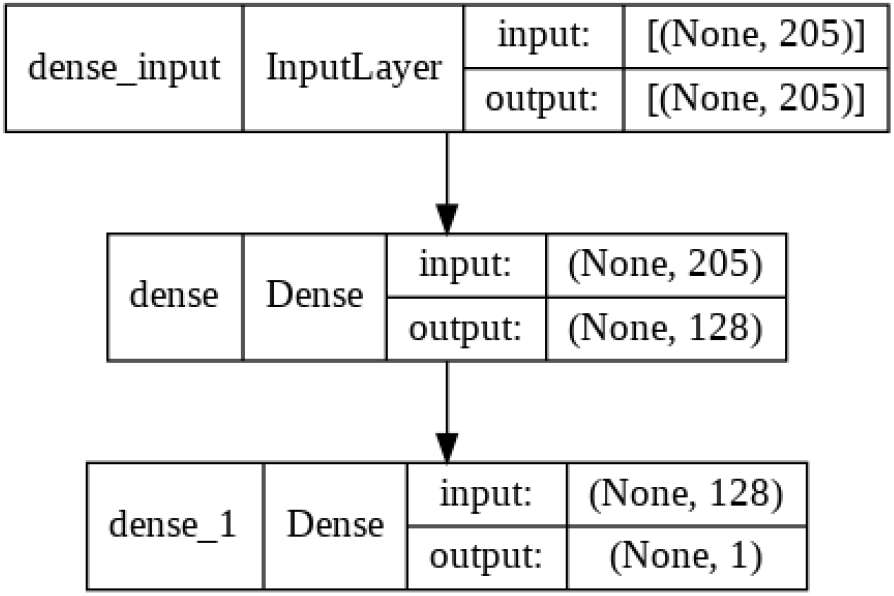
Proposed Model Architecture For Section 3.4.1 and 3.4.2

#### 3.4.2 Combining 100 Balanced Data-set and Training a Model

The aforementioned 100 balanced sampled datasets are combined to obtain a single dataset. Then, we trained and evaluated the same model that we used for the imbalanced dataset.

#### 3.4.3 Multi- and Single-Input Ensemble Model

As a third approach, we build a multi-input ensemble model. As shown in Figure 5, 10 identical models are combined right before their output layers and forced to yield a single output. The advantage of this approach is that models are trained as a group. Therefore, the weights on the concatenation layer are adjusted while training. Besides, initial weights are different for each of the 10 models since they are generated randomly before the training phase.

**Fig. 5:**
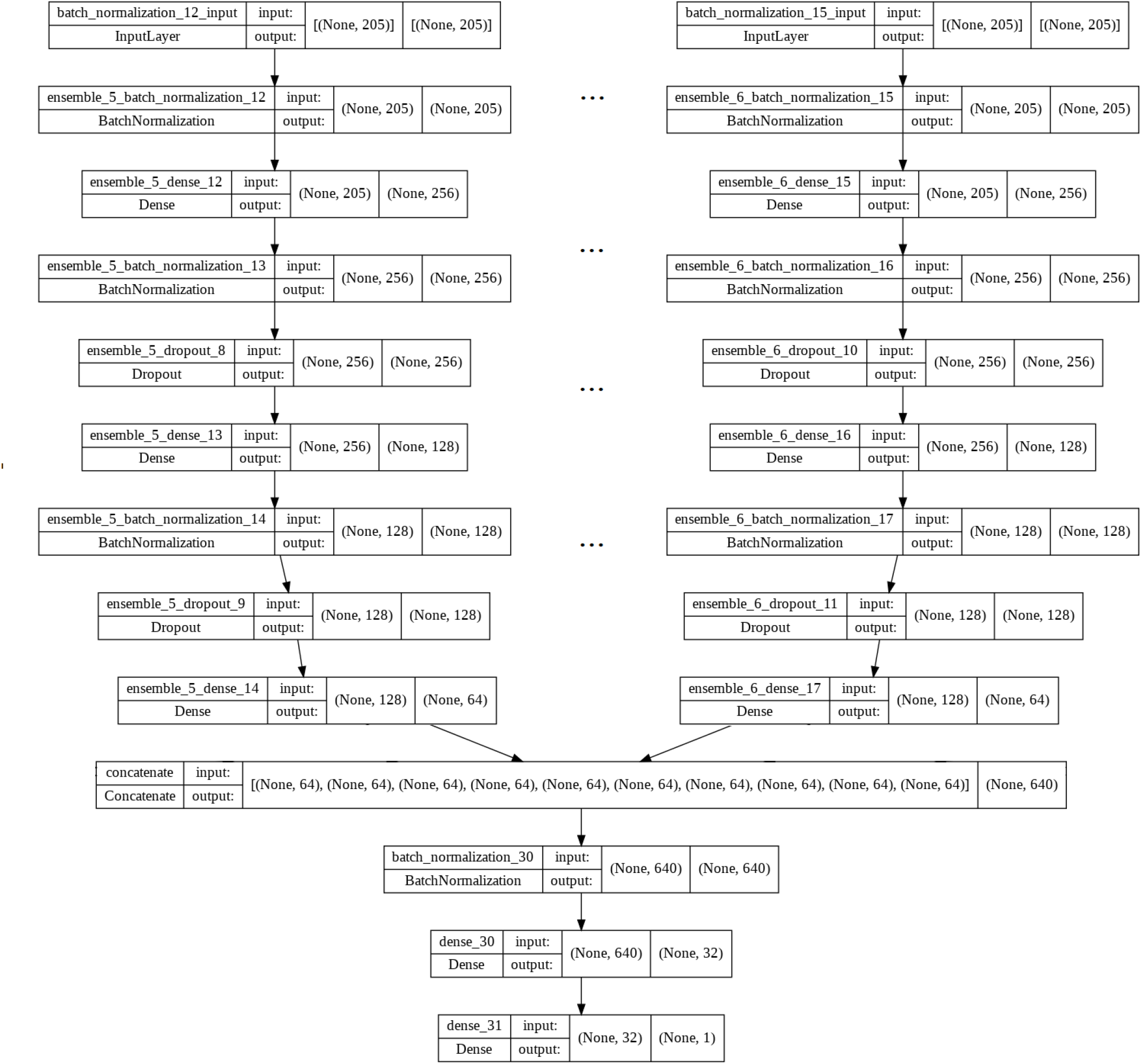
Proposed Model Architecture For Multi-Input Ensemble Model

In particular, the model architecture consists of two parts: a head part (before the concatenation layer) and a tail part (after the concatenation layer). In the head part, we have 10 models that share the same architecture and are trained with the same dataset. The key difference here is that each model is generated with random initial weights that affect the learning phase of the model slightly. In the model, we employ fully connected dense layers, Batch Normalization [41] is exploited to normalize the output of the previous layer, and Dropout [42] is used to avoid the over-fitting problem.

The first layer in the model is the batch normalization layer which normalizes the input features. Then, we create a block of layers that includes a fully connected dense layer, a batch normalization layer, and a dropout layer. We repeat that block two times sequentially. In the first block, a fully connected dense layer extends 205 input values to higher dimensions with 256 units. A normalization layer normalizes the output of the dense layer and the dropout layer regularizes the units with a 0.5 ratio. In the second block, we downsize the fully connected layer units from 256 to 128. Similar to the first block a batch normalization layer and a dropout layer with a 0.3 regularization ratio follow it. Concatenating the outputs of 10 models would be enormous with 128 for each. Therefore, we add one more layer that has 64 units right after the sequential blocks to reduce the cost of the concatenation operation.

The concatenation layer outputs 640 features as expected. In the tail part, we normalize 640 units with a batch normalization layer and a dense layer that downsizes 640 features to 32 features. To predict the class of the samples, an output layer with a sigmoid activation function is used.

Multi-input model is costly and takes relatively much more time to train for each epoch. Hence, we combine the inputs of the models and modify the baseline model architecture slightly to use as head part of the model. As a result, we build a new ensemble model that has a single input and single output. Figure 6 visualizes the architecture of the new single-input ensemble model. In this case, we keep the parameters of the head part relatively small, but the concatenation layer has a high dimension when compared to the multi-input model. Similar to the multi-input model, we apply batch normalization to the input features. However, the difference is that right after the batch normalization layer, we downsize the features from 205 to 128, concatenate 10 different 128 units, and finally, apply L2 regularization with the ratio of 0.01 instead of Dropout. Since we doubled the size of the concatenation layer, a dropout layer with a 0.8 ratio is added after the concatenation layer to regularize the model. A fully connected layer that has 32 units and L1 regularization with a 0.011 ratio follows the dropout layer. Then, the model ends with a second dropout and an output layer.

**Fig. 6:**
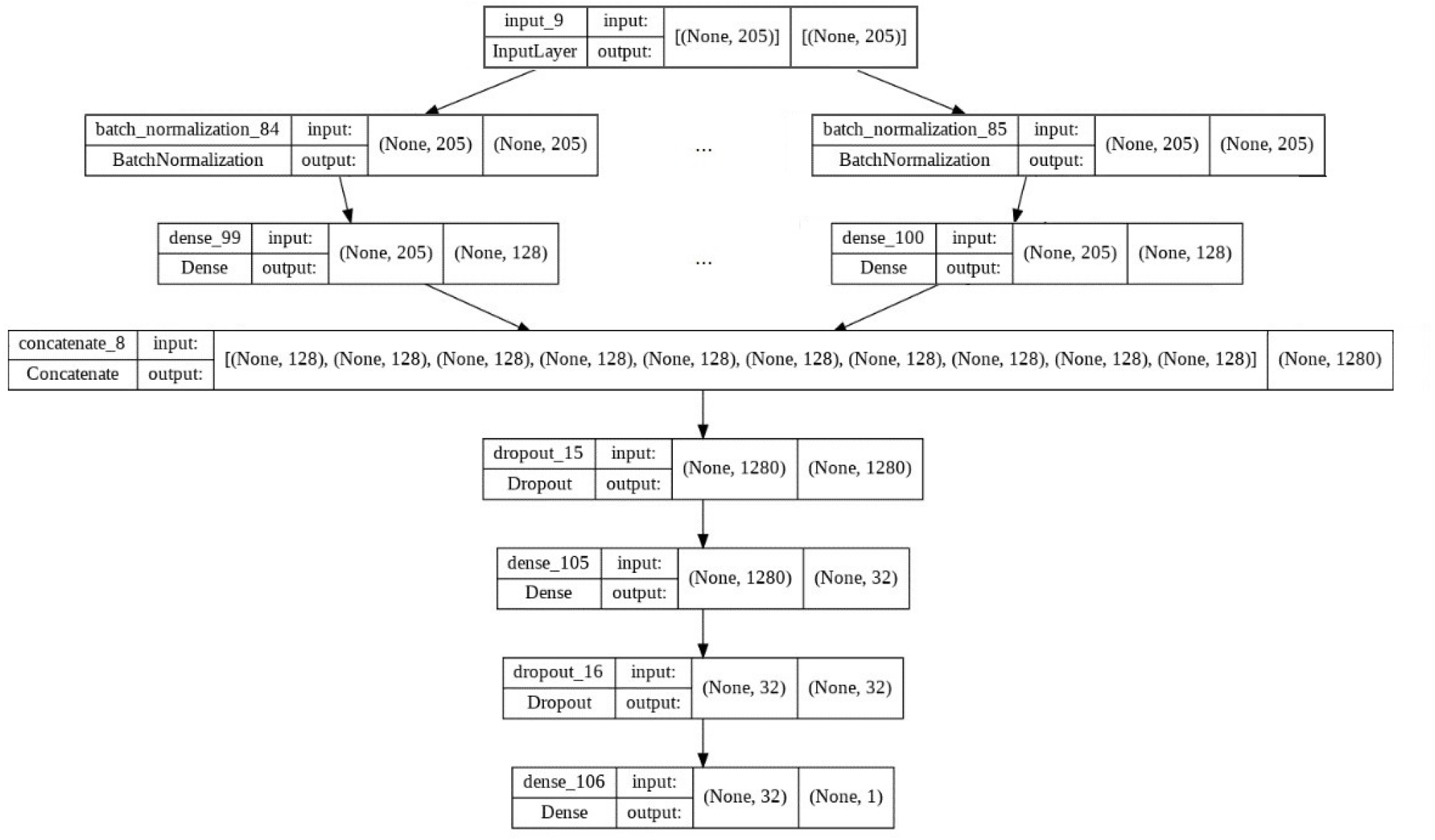
Proposed Model Architecture For Single-Input Ensemble Model

In brief, we apply both Dropout and L1-L2 regularization to the model. That allows us to train the model with more epochs (e.g., 16) without much over-fitting.

#### 3.4.4 Bagging-Classifier With 100 Model

In this approach, a Bagging classifier [43] is built with 100 neural network models. In particular, each model is trained using a different sampled dataset. Therefore, instead of looking at the test samples from the same perspective, each model would see them from a different point of view. After training each model separately, we apply hard and soft voting [44] to compare the results. In hard voting, regardless of the confidence levels of models, the most voted label is selected as the final decision. However, in soft voting, the confidence levels of models are considered and the average of label percentages is considered while determining the final label for a sample. The proposed model architecture is shown in Figure 7.

**Fig. 7:**
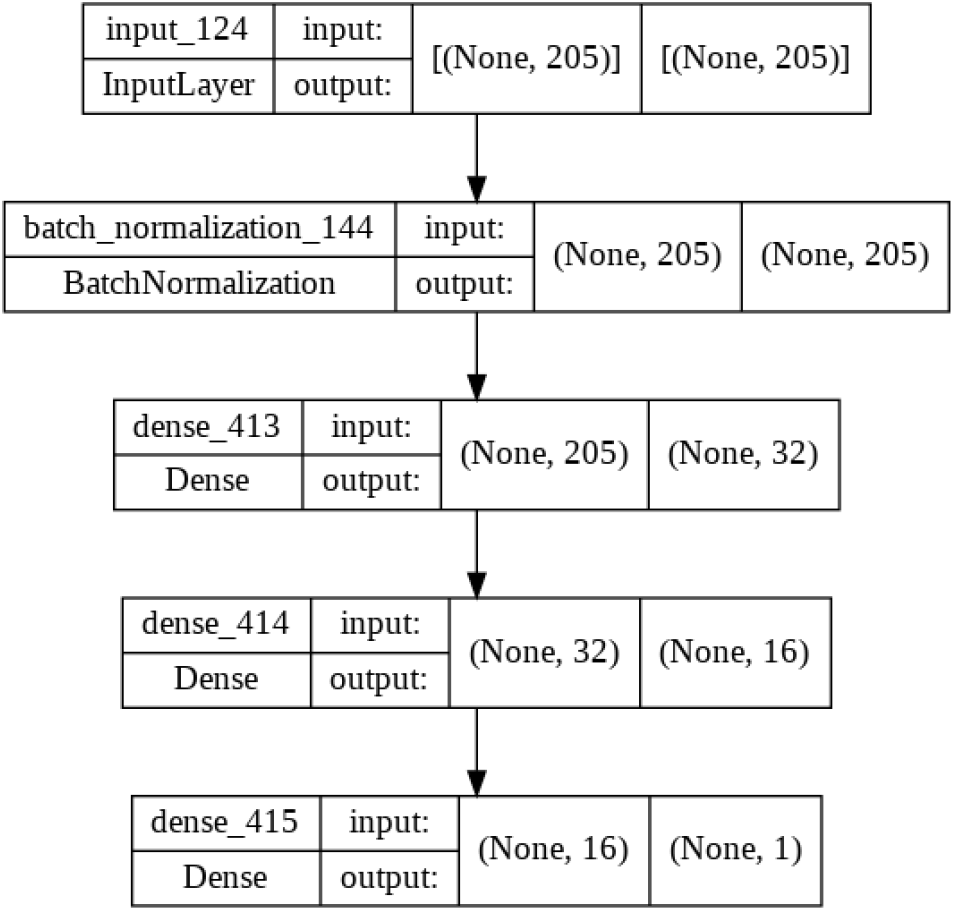
A Model Architecture For Bagging Classifier

This model is also a slightly modified version of the baseline model. In particular, instead of downsizing the feature sizes from 205 to 128, in this case, we reduce the feature sizes to 32. Then, we insert a new 16 units layer. Then, the output layer follows it.

#### 3.4.5 Training an NLP Model For Seq-2-Seq Generation

T5 [45] is a Natural Language Processing (NLP) model pre-trained on a multi-task mixture of unsupervised and supervised tasks. The difference between T5, Bert [46], which has only encoder blocks, and GPT-2 [47], which has only decoder blocks, is that T5 combines both Bert and GPT-2. Thanks to this feature, researchers have trained many models using T5 models in different tasks like language translation. In our case, we trained the model to generate mutated sequences from the non-mutated sequences. More specifically, the model is trained with different coding regions (‘E’, ‘M’, ‘N’, ‘ORF10’, ‘ORF1ab’, ‘ORF3a’, ‘ORF6’, ‘ORF7a’, ‘ORF7b’, ‘ORF8’, ‘S’). In order to obtain proteins, sequences are converted from base pairs to amino acids using the NCBI website. Then, each protein region is separated from a sequence using their corresponding start and end points and each region is considered as a different data point. Fig. 8 shows the length of ORF1ab as an example of the protein lengths.

**Fig. 8:**
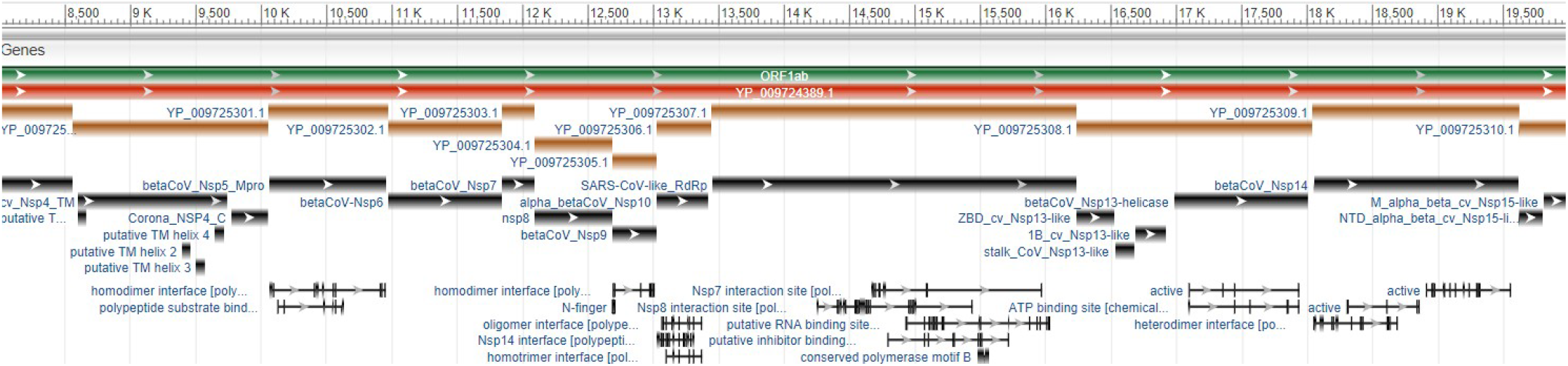
ORF1ab Sequence Length: ScreenShot from NCBI website

## 4 Results

In this section, the results of each approach and model are evaluated in detail with pros and cons. Evaluating models with a single metric would be misleading for unbalanced and even for balanced datasets. For instance, let’s say we have 900 samples for the non-mutated class and 100 samples for the mutated class. The model would get 90% accuracy even if it predicts all samples as non-mutated. Therefore, we evaluated models with various metrics such as precision, recall, f1-macro, etc. Lastly, Positive Predictive Ratio (PPR) and Negative Predictive Ratio (NPR) [48] are used as additional metrics.

In addition, for the NLP approach, we propose a new metric called *PAM metric*. The name comes from the PAM250 matrix [49]. In particular, we utilize the PAM250 matrix of the NCBI in which the entries are at a scale of *ln*(2)*/*3. In the matrix, each amino acid has different scores for mutating into other amino acids or itself (i.e., staying the same). For instance, if “A” mutates to “A”, the corresponding score for this mutation would be 2 which we call the *true value*, but if amino-acid “A” mutates to “R”, the score would be -2. Based on these mutation scores, we locate the score of each amino-acid change within a predicted sequence and sum up their scores. Then, the calculated score is divided by the true value of the sequence for normalization purposes. As an example, assume that the actual (i.e., true) sequence consists of amino acids “A A B B C” and our model’s prediction is “A A C B”. Note that the predicted sequence is missing one amino acid. For these cases, we fill * symbol for each missing amino acid in order to penalize the gaps. Thus, we end up with the prediction sequence “A A C B *”. In the evaluation phase, the true value before the prediction would be the summation of 2+2+3+3+12 for “A A B B C” which sums up to 22 and the model’s prediction score would be the summation of 2+2+(-4)+3+(-8) for sequence “A A C B *” which sums up to -5. After normalization, the final result would be *Value*_*predicted*_*/Value*_*true*_ = −5*/*22 = −0.227. In the proposed PAM-based metric, the best score is 1.0 for a perfect prediction and the worst score is negative infinity. However, as a ground truth, we expect the model to perform at least the score of the original sequence which does not contain the mutations that we aim to predict.

Nevertheless, considering a single metric might be misleading for long sequences. For instance, if there are a few mutations within an amino acid sequence, a model may predict the whole sequence the same as the original to easily achieve a score of 0.99. To eliminate these cases and obtain a fair evaluation, we extract all mutated amino acids from a sequence and generate a new sequence that only consists of the mutated amino acids. Then, we apply the same scoring procedure to evaluate a model performance to predict the mutated amino acids.

For all the presented confusion matrices in this section, 1 denotes the mutated samples and 0 denotes the non-mutated samples.

### 4.1 Baseline: Model With Imbalanced Data

As shown in Fig. 9, the model is biased to predict the majority class label which is 0 (non-mutated) for test samples since the data is quite imbalanced.

**Fig. 9:**
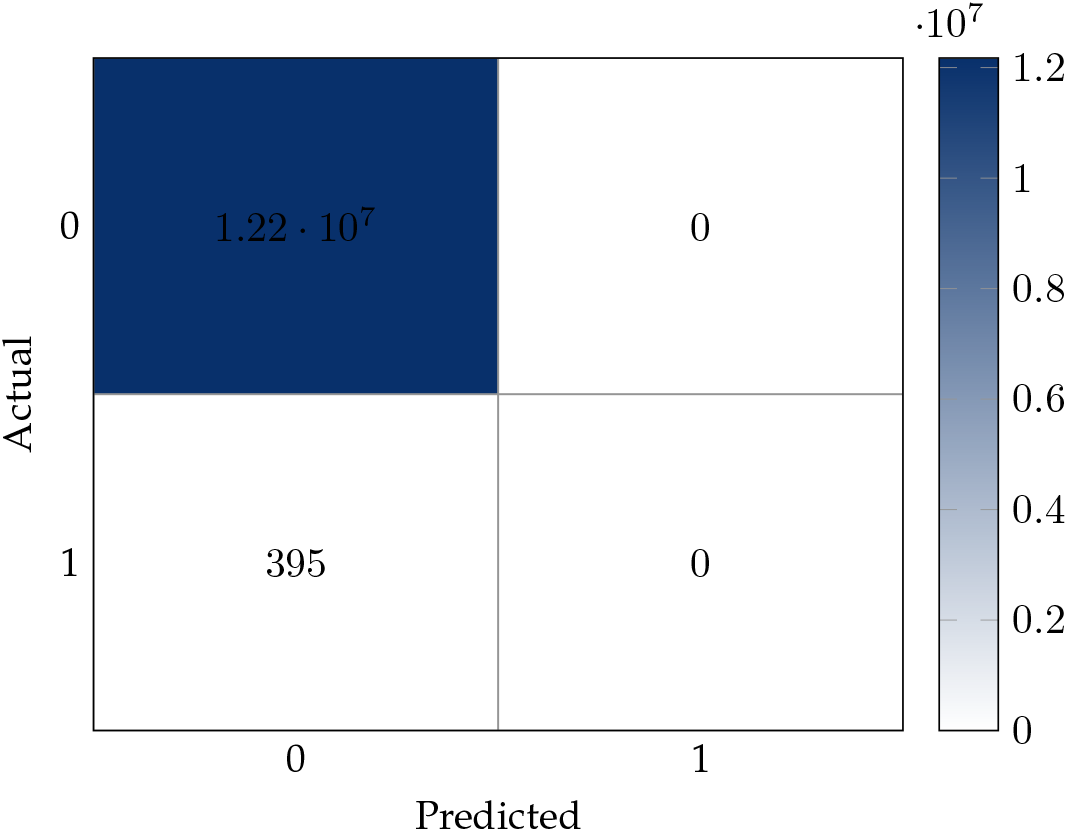
Test results of the model trained with the imbalanced dataset: Confusion matrix

### 4.2 Combining 100 Balanced Data-set

In this approach, 100 sampled balanced datasets are combined and fed into a single model. The performance of the model is presented in Fig. 10. On the test dataset, a single model predicts the non-mutated and mutated samples well as shown by the confusion matrix.

**Fig. 10:**
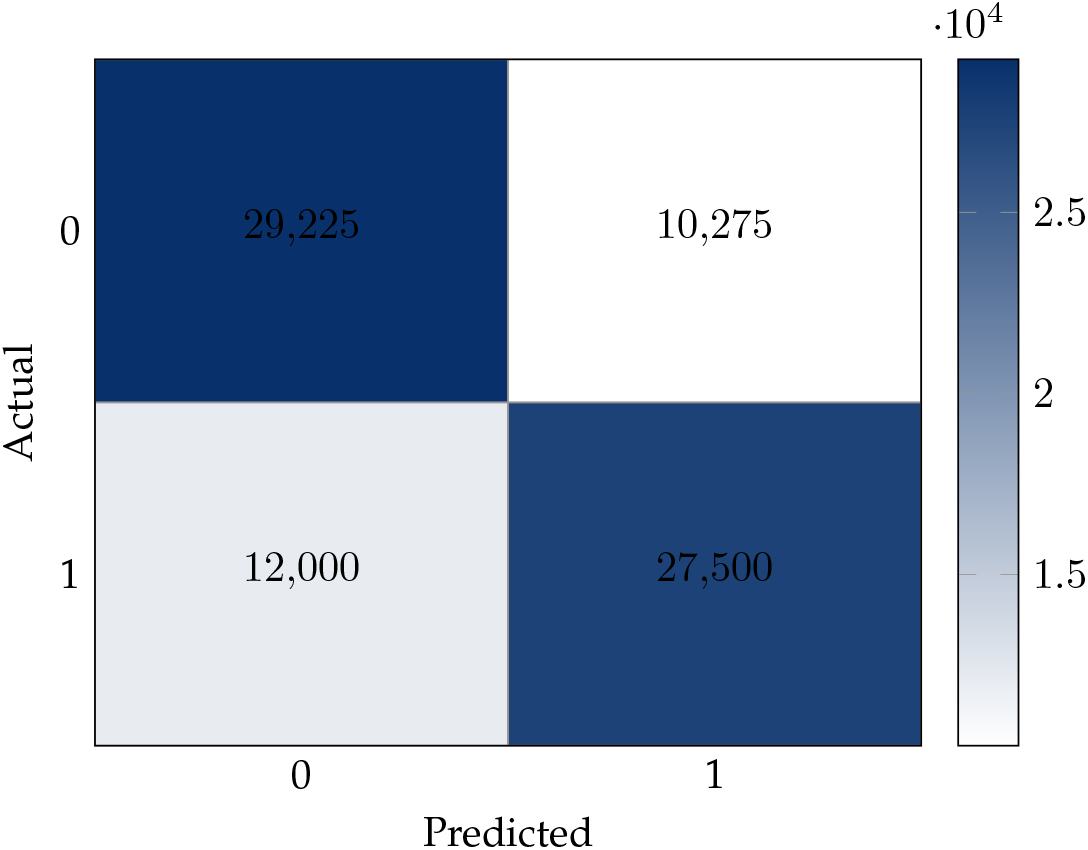
Test results of the model trained with the balanced dataset: Confusion matrix

### 4.3 Multi- and Single-Input Ensemble Model

In order to obtain a better score on the combined dataset, we created a multi- and single-input ensemble models and the performance results are shared in Figures 11 and 12.

**Fig. 11:**
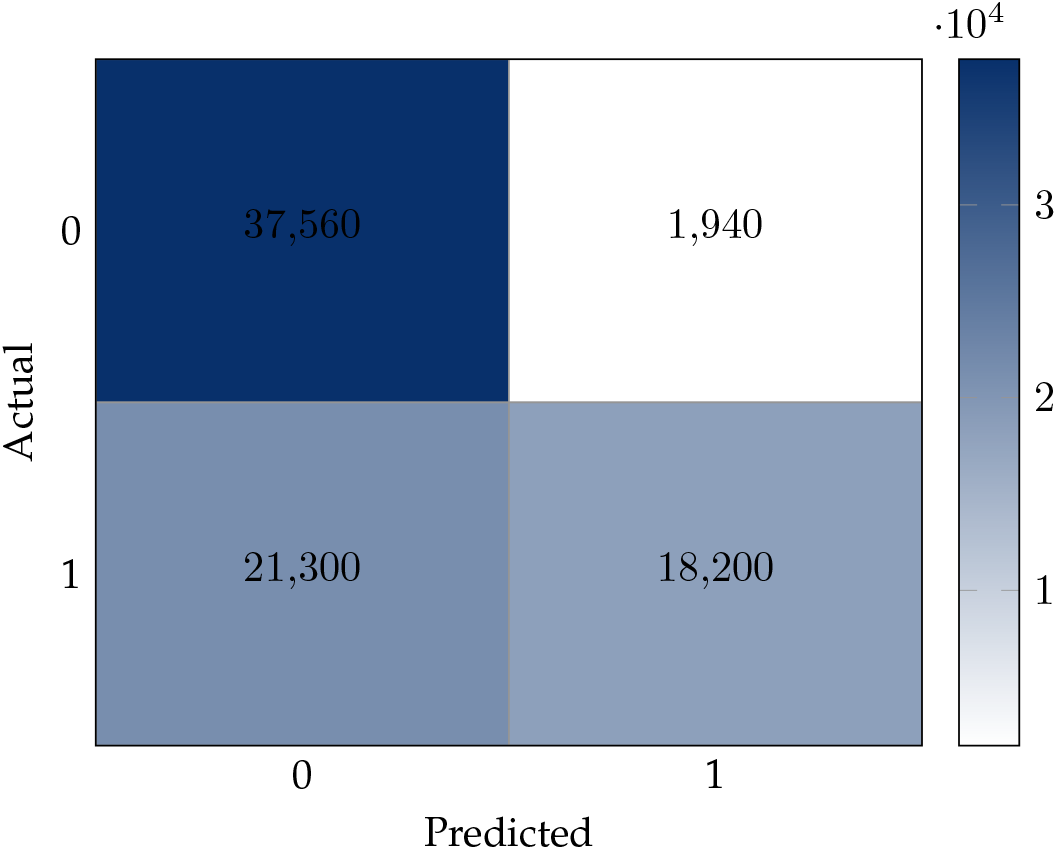
Test results of the Multi-Input Ensemble Model: Confusion matrix

**Fig. 12:**
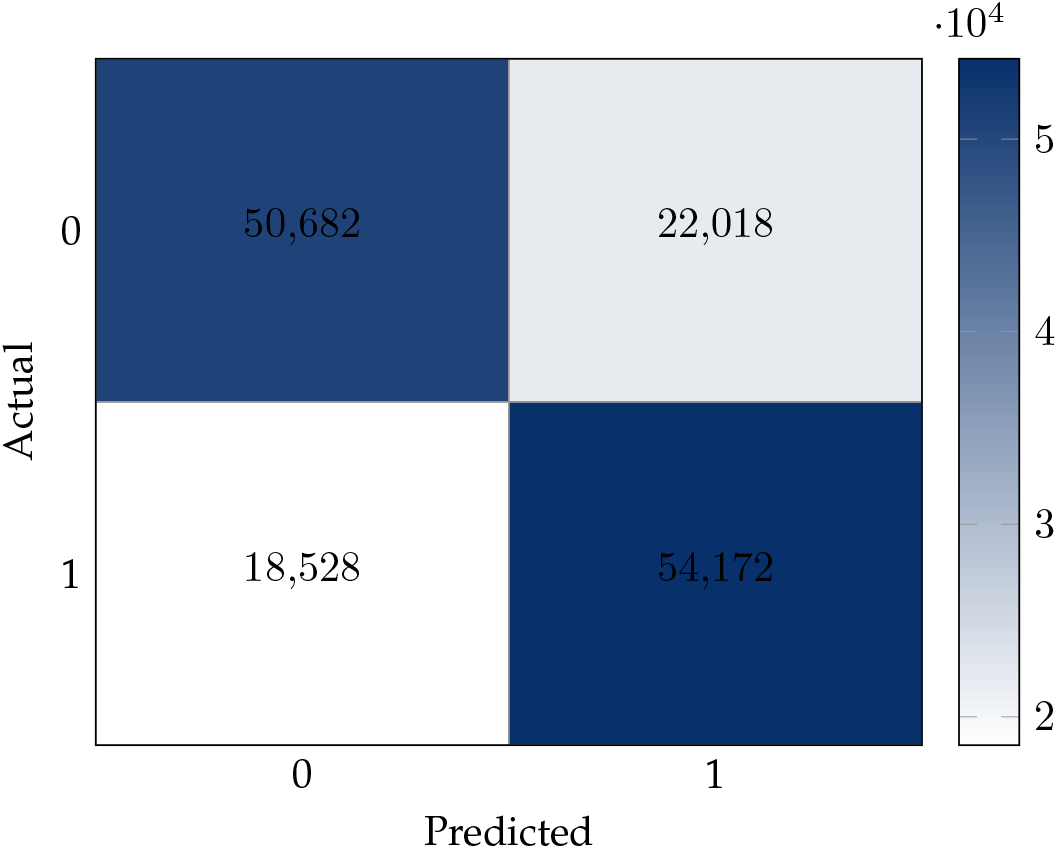
Test results of the Single-Input Ensemble Model: Confusion matrix

The multi-input ensemble model tends to predict more than half of the mutated samples as non-mutated and the performance of it is not quite well in terms of recall of the mutated samples and overall f1-macro score. That means, even though the model can classify most of the non-mutated samples as correct (95% recall), it classifies mutated samples poorly (46% recall). On the other hand, the performance of the single-input ensemble model is promising in terms of all the considered metrics.

### 4.4 Bagging-Classifier With 100 Model

To build a bagging classifier, 100 models each of which has the same architecture and is trained with different balanced datasets. Figure 13 shows the distribution of mutation percentages for all predictions made by 100 models for each sample. Since the employed threshold for positive class prediction is 0.5, the percentages of the predictions which is close to 1.0 have a higher confidence level for the class label “mutated” (i.e., 1). Likewise, percentages close to 0 have a higher confidence level for the class label “non-mutated” (i.e., 0).

**Fig. 13:**
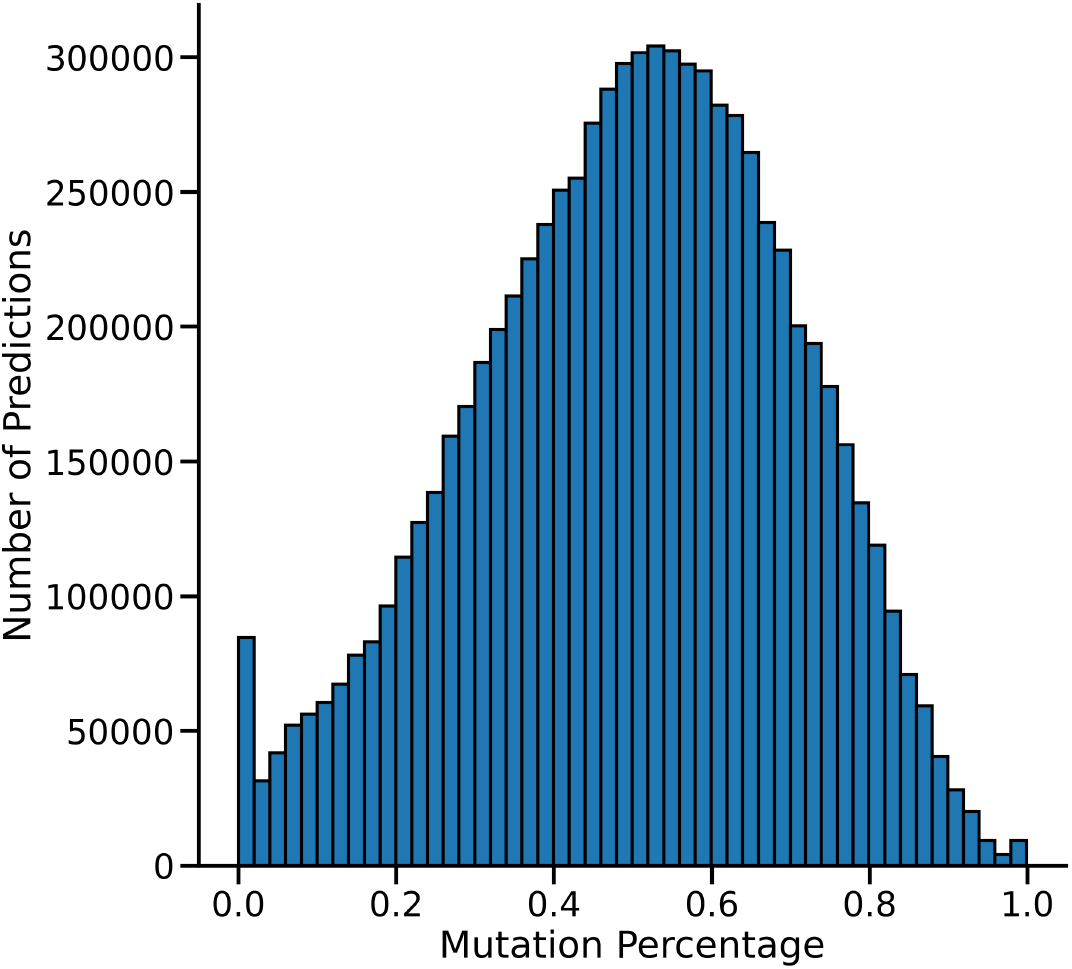
Distribution of the prediction of 100 Models

In this case, since we have multiple models, we apply hard-voting method which is also called majority voting. Considers the prediction of all models to make the final decision about the mutation status of a position in the genome based on the majority vote. Similar to the previous cases, for each model we set a threshold of 0.5%, and consider those below the threshold as non-mutated while those above the threshold as mutated. Fig. 14 shows the performance of the final model.

**Fig. 14:**
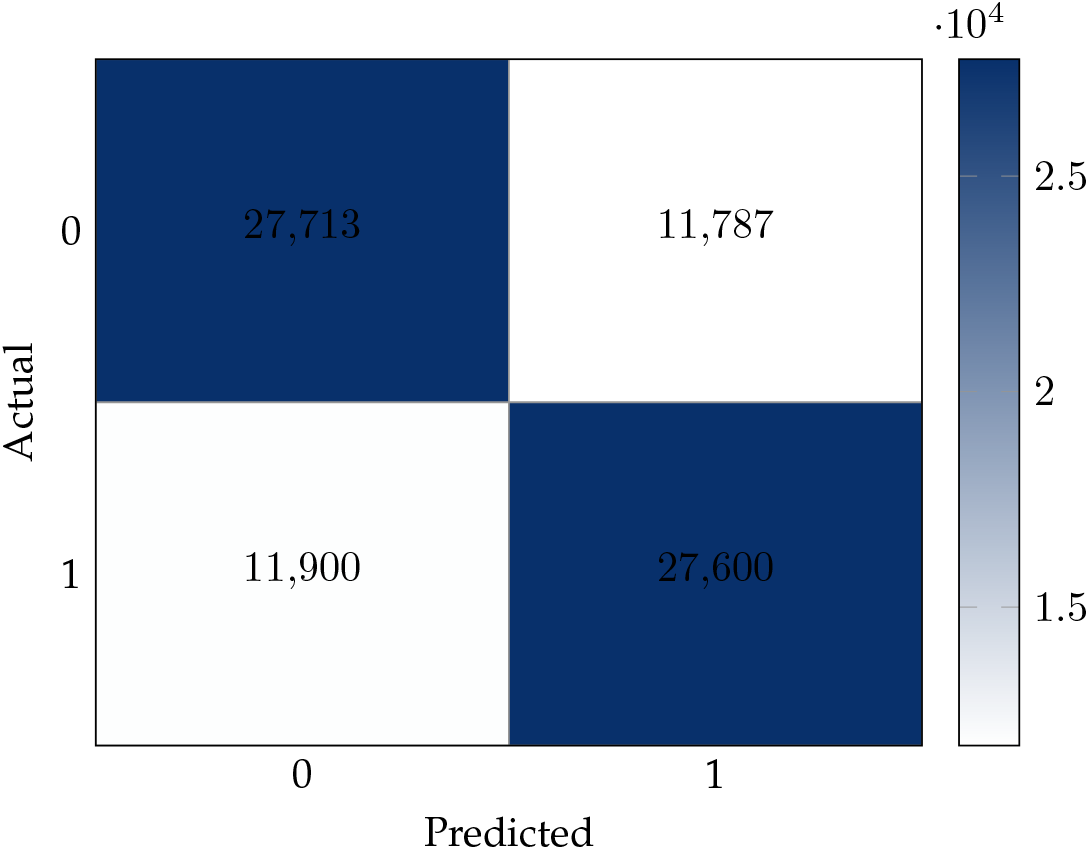
Test results of the Bagging Classifier: Confusion matrix

### 4.5 Training an NLP Model For Seq-2-Seq Generation

Seq-2-Seq is an NLP task that is quite different from the previous machine learning approaches. For this approach, we generated two different metrics to evaluate the trained model. In terms of PAM score-based metric, we obtain 99% accuracy. That means the model can generate child sequences very well. However, if we look at the result of the PAM score-based metric, the model is missing almost all mutated amino acids. The original score, which is a base score before the mutation happens, is roughly -0.45. However, after the prediction, the score is roughly -0.44, which is slightly better than the original score. Thus, we can consider it as missing almost all mutations since the score of the prediction is almost same with the score of parent sequence which is the score before mutation. A similar seequence-to-sequence generation method was recently employed by Mohamed et al. [9] to predict mutations in Influenza genome. The results from this comparison demonstrate that new approaches are required for mutation prediction in SARS-CoV-2 genome.

### 4.6 Putting It Altogether: A Comparison of All Models

Fig. 15 compares all the proposed models in a single chart in terms of their F1 scores. To sum up, among various models that we consider in this paper, the Single-Input Ensemble Model and Bagging-Classifier yield the best scores.

**Fig. 15:**
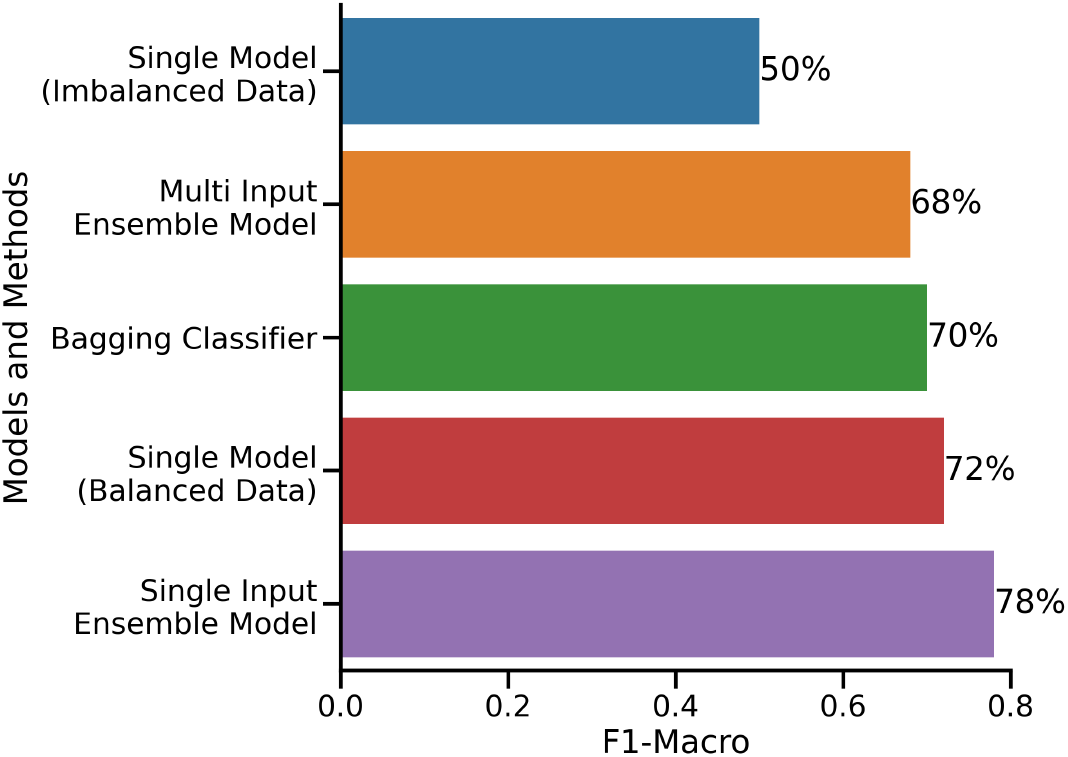
Comparing the F1-Macro Scores of All Methods

## 5 Conclusion

SARS-CoV-2 challenged the humanity in so many different ways from the dramatic disruption of people’s social and professional lives to the incapacity of the health systems in responding to the enormous need for intensive care demand of the patients. Even though several vaccines were developed to help stop the pandemic, rapid mutation rates of the virus poses great threat to the efficacy of the vaccines. Thus, foreseeing the mutations well before they emerge may help future planning future vaccine designs. To this end, in this study, we propose an evaluate a variety of deep learning-based machine learning models to predict the future mutations of the SARS-CoV-2 genome. Our results show that single input ensemble and bagging-classifier models perform the best with an F1 score of 0.78. We hope that this study will contribute getting ahead in the fight against SARS-CoV-2 and similar future viruses that may arise in the future.

This work may be extended in several different ways. First, the effect of each possible mutation on the corresponding protein structure may be computed. Then, the distance between the original and the mutated structure of each protein may be included as additional features in prediction models. Second, on a high performance computing cluster, the whole dataset may be used to train the models. More data may lead to higher accuracy models. Third, based on mutation profiles, the predicted future strains of SARS-CoV-2 may be assesed in terms of their virulence. In particular, the statistical models proposed in this work may be adapted for such predictions. This would enable detecting potentially dangerous strains ahead of time, and getting prepared for it earlier to reduce the impact.

## 6 Acknowledgments

We thank Dr. Ogun Adebali and Cem Azgari for providing the mutation profile data.

## 7 Declarations

### 7.1 Ethics approval and consent to participate

Not applicable.

### 7.2 Consent for publication

Not applicable.

### 7.3 Availability of data and materials

Data is available on GISAID portal for all researchers. Source codes are available upon request from the authors.

### 7.6 Authors’ contributions

HA and ARI contributed to the writing of the manuscript, implemented the data processing, model training, and evaluation. AA prepared the GISAID data for model processing and implemented the pipelines for building mutation profiles. MB is involved in model development and evaluation. MEB conceived the problem and supervised the genomic side of the research. GD is involved in model development and evaluation. EK helped Swith the data processing and manuscript writing. BO contributed to the feature identification for models. AC acquired funding, contributed to the writing of the manuscript, and supervised the overall research.

